# Functional Characterization of SARS-CoV-2 3CLpro Variants

**DOI:** 10.64898/2025.12.23.696280

**Authors:** Dipon Saha, S. Yasin Tabatabaei Dakhili, Eliot Mar, Yu Seby Chen, Filip Van Petegem, Dieter Brömme

**Author notes:** Authors contributed equally (Dipon Saha, S. Yasin Tabatabaei Dakhili).

## Abstract

The SARS-CoV-2 main protease (3CLpro) is essential for viral replication and a leading antiviral target. Circulating variants accumulate substitutions on this enzyme, distant from the catalytic site. Surprisingly, mutant enzymes retain full proteolytic activity, though preserved overall activity does not exclude subtler effects on substrate recognition or selectivity, arising from distal structural perturbations. In this study, we compared the steady-state kinetics of wild-type (Wuhan) 3CLpro with enzymes from the Beta (K90R), Lambda (G15S), and Omicron (P132H) variants, using two peptide substrates representing distinct viral polyprotein cleavage sites. All four proteases displayed comparable catalytic efficiencies, similar pH-rate profiles, suggesting conservation of the catalytic mechanism despite sequence variation. The crystal structure of Omicron 3CLpro bound to an Nsp8-Nsp9 peptide revealed a conserved fold and active-site geometry, with the P132H side chain adopting a substrate-dependent conformation that rebuilt its local contacts – indicating towards how a distal substitution is accommodated without perturbing catalysis. Thermal stability measurements identified the sole distinguishing effect of P132H, with Omicron showing altered stability at elevated temperature. A screen of 31 tanshinones against 3CLpro identified T06 with K_i_ values of 5 µM, as 3CL pro inhibitor. Thus, 3CLpro may tolerate distal substitutions through local structural adaptation, supporting its durability as an antiviral target.

## Introduction

The COVID-19 pandemic has made clear the urgent need for antiviral strategies that remain effective as SARS-CoV-2 continues to evolve^1,2^. While vaccines primarily target the spike protein and are vulnerable to antigenic drift, antiviral drugs directed against conserved viral enzymes offer a complementary approach that may retain efficacy across variants^3,4^. Among these targets, the main protease, 3-chymotrypsin-like protease (3CLpro) is an attractive therapeutic target, exemplified by the clinical inhibitor Nirmatrelvir. 3CLpro cleaves the SARS-CoV-2 polyproteins (pp1a and pp1ab) at eleven distinct sites to release functional non-structural proteins (Nsp)^5,6,7^. This functional importance makes it necessary to assess how variant-associated amino acid substitutions in 3CLpro may alter enzymatic function that could compromise therapeutic efficacy^8,9,10,11,12^.

To date, experimental characterization of 3CLpro has focused largely on the Nsp4-Nsp5 (`AVLQ`↓`SGFR`) cleavage site, even though the protease must process ten additional junctions that differ in sequence composition and physicochemical properties^13,14,15^. This limited substrate scope leaves unresolved whether variant-associated mutations influence 3CLpro function across cleavage sites with differing sequence composition.

A particularly informative test case is the Nsp8-Nsp9 junction, which contains a P′-site composition featuring two consecutive asparagine residues (P1’ and P2’) that may impose distinct structural and catalytic constraints^13,14,16,15^. The impact of variant-dependent mutations on this cleavage site has not been systematically assessed, and structural information on how variant enzymes engage this substrate is still lacking. These gaps raise the possibility that mutations distant from the catalytic residues could produce functional impact that are not apparent when only the Nsp4-Nsp5 site is examined.

Given the need for inhibitors that retain activity against variant proteases, compounds with alternative binding mechanisms and established anti-coronaviral activity warrant further investigation. In this context, tanshinones have emerged as antiviral candidates. These diterpenoid quinones, derived from *Salvia miltiorrhiza*, possess diverse pharmacological activities, including anti-inflammatory, antioxidant, and enzyme-inhibitory properties^17^. Several members of this compound family have been reported to inhibit coronavirus proteases, including SARS-CoV and SARS-CoV-2 3CLpro^18,19,4^. Unlike covalent inhibitors such as Nirmatrelvir, tanshinones may act through both non-covalent and covalent mechanisms, providing alternative binding modes that could be less vulnerable to resistance arising from active-site mutations^20,21^. Despite these characteristics, their efficacy against SARS-CoV-2 variants and their ability to maintain inhibitory activity in the presence of naturally occurring 3CLpro substitutions remain poorly understood. Therefore, systematic evaluation of tanshinones across variant proteases is needed to define their potential as broad-spectrum inhibitors and to assess their utility in expanding the antiviral arsenal against evolving SARS-CoV-2 lineages.

The present study provides a comparative functional analysis of WT-3CLpro and the 3CLpro variants of other SARS-CoV-2 lineages (Beta, Lambda, and Omicron) to determine how sequence variation influences activity across distinct canonical cleavage sites, substrate processing, and inhibitor responses. We characterized enzymatic properties using the widely studied Nsp4-Nsp5 substrate and extended the analysis to the structurally distinct Nsp8-Nsp9 junction to assess substrate-dependent effects. We also determined the crystal structure of the Nsp8-Nsp9 peptide bound to the enzyme. In parallel, we examined the pH and thermal profiles of these variant enzymes. We further evaluated susceptibility to the clinical inhibitor Nirmatrelvir and screened a panel of tanshinones to probe the potential of alternative inhibitors. Together, these studies address whether distal mutations in 3CLpro alter catalytic function, substrate recognition, or inhibitor sensitivity, providing insight into the resilience and adaptability of this essential viral enzyme as SARS-CoV-2 continues to diversify.

## Results

### Steady-State Kinetics of WT and Variant 3CLpro Enzymes

To evaluate the impact of single amino acid substitutions on the proteolytic activity of 3CLpro variants, we measured steady-state kinetic parameters using a Förster resonance energy transfer (FRET) substrate, (MCA)`AVL`Q↓`SGFR`[`K`(Dnp)]`RR`, which corresponds to the native Nsp4-Nsp5 cleavage site. Here, we analyzed 3CLpro from WT (Wuhan), Beta (K90R), Lambda (G15S), and Omicron (P132H) lineage. All these mutations are ∼17-20 Å away from the active site (Fig. 1A, B). PISA^22^ analysis of X-ray crystal structures of WT SARS-CoV-2 3CLpro (e.g. PDB: 7JP1)^15^, indicate that the residues G15, K90 and P132 are solvent-exposed. Except for the mutations P132H, the other mutations introduce no major changes in the chemical character of the sidechains. The mutations G15S and K90R are in domain I, whereas the mutation P132H is in domain II. To the best of our knowledge, these residues participate neither in the active site nor the allosteric binding sites of SARS-CoV-2 3CLpro.

**Figure 1:**
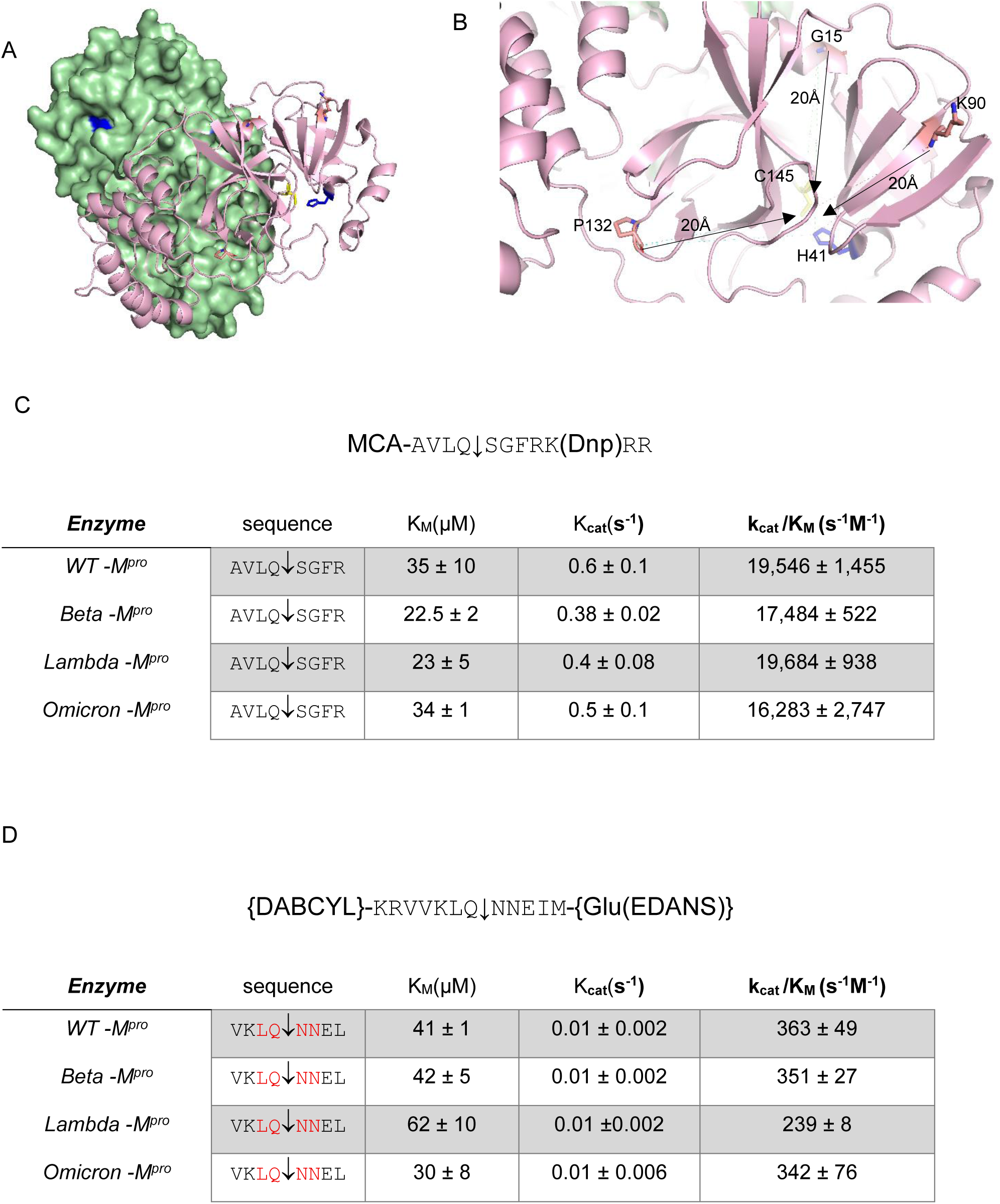
Kinetic analysis of 3CLpro from various SARS-CoV-2 variants in the presence of two substrates representing the Nsp4-Nsp5 and Nsp8-Nsp9 processing sites. A) X-ray crystal structure of SARS-CoV-2 3CL-Mpro (7JP1) indicating the catalytic dyad (in stick model; blue and yellow) and key mutation (stick model) B) Zoom in view of the sites of mutations (stick model) and the catalytic dyad (blue and yellow) are indicated. C) Table compares the kinetic parameters of 3CL-Mpro using substrate MCA-`AVLQ`↓`SGFRK`(Dnp)`RR`. D) Table compares the kinetic parameters of 3CL-Mpro using a substrate ({DABCYL}-`KRVVKLQ`↓`NNEIM`-{Glu(EDANS)}).

Recombinant WT-3CLpro exhibited a catalytic efficiency (k_cat_/K_M_) of 19,546 ± 1,455 M^−1^ s^−1^ (K_M_ = 35 ± 10 µM; k_cat_ = 0.6 ± 0.1 s^−1^). Variant enzymes, including Omicron (P132H), Beta (K90R), and Lambda (G15S), display catalytic efficiencies within the same order of magnitude as WT (Fig. 1C). For example, the Omicron variant had a k_cat_/K_M_ of 16,283 ± 2,747 M^−1^ s^−1^ (K_M_ = 34 ± 1 µM; k_cat_ = 0.5 ± 0.1 s^−1^), indicating similar kinetic parameters under these assay conditions (Fig. 1C)^10,11,12^

We performed the kinetic analysis using an alternate FRET substrate that mimics the Nsp8-Nsp9 cleavage site, (DABCYL)`KRVVKLQ`↓`NNEIME`(EDANS). Catalytic efficiencies for this substrate were lower, by up to eightfold, compared with the Nsp4-Nsp5 substrate, yet the Omicron variant (342 ± 76 M^−1^ s^−1^) remained similar to WT-3CLpro (363 ± 49 M^−1^ s^−1^) (Fig. 1D)^13,14^. Beta and Lambda 3CL-Mpro have shown similar efficiency in processing this substrate (350 ± 27 M^−1^ s^−1^ and 239 ± 7 M^−1^ s^−1^).

Collectively, these data indicate that naturally occurring variant-associated substitutions located distal to the catalytic site do not measurably alter the intrinsic proteolytic efficiency of 3CLpro toward either the canonical Nsp4-Nsp5 junction or the alternative Nsp8-Nsp9 cleavage site, supporting the conclusion that the catalytic function of the protease remains conserved across major SARS-CoV-2 lineages.

### Co-crystal Structure of Omicron 3CLpro Bound to an Nsp8-Nsp9 Cleavage-Site Peptide

Although steady-state kinetics showed that the P132H Omicron variant retains WT-like catalytic efficiency, these measurements do not reveal whether this preserved activity reflects an unchanged catalytic architecture or compensatory structural rearrangements that may influence substrate recognition and inhibitor susceptibility despite similar overall turnover. To define the molecular basis of this preserved activity and assess whether the distal P132H substitution induces long-range conformational effects on substrate recognition, we determined the co-crystal structure of Omicron 3CLpro bound to a peptide derived from the Nsp8-Nsp9 cleavage sequence (Fig. 2). Structures were resolved to 1.94 Å (Table S1). The C145A substitution was introduced to trap the processed substrate (Fig. 2A). Because the C145A substitution removes the catalytic nucleophile, the peptide is captured before cleavage, so the structure represents the Michaelis complex rather than an acyl-enzyme intermediate.

**Figure 2.**
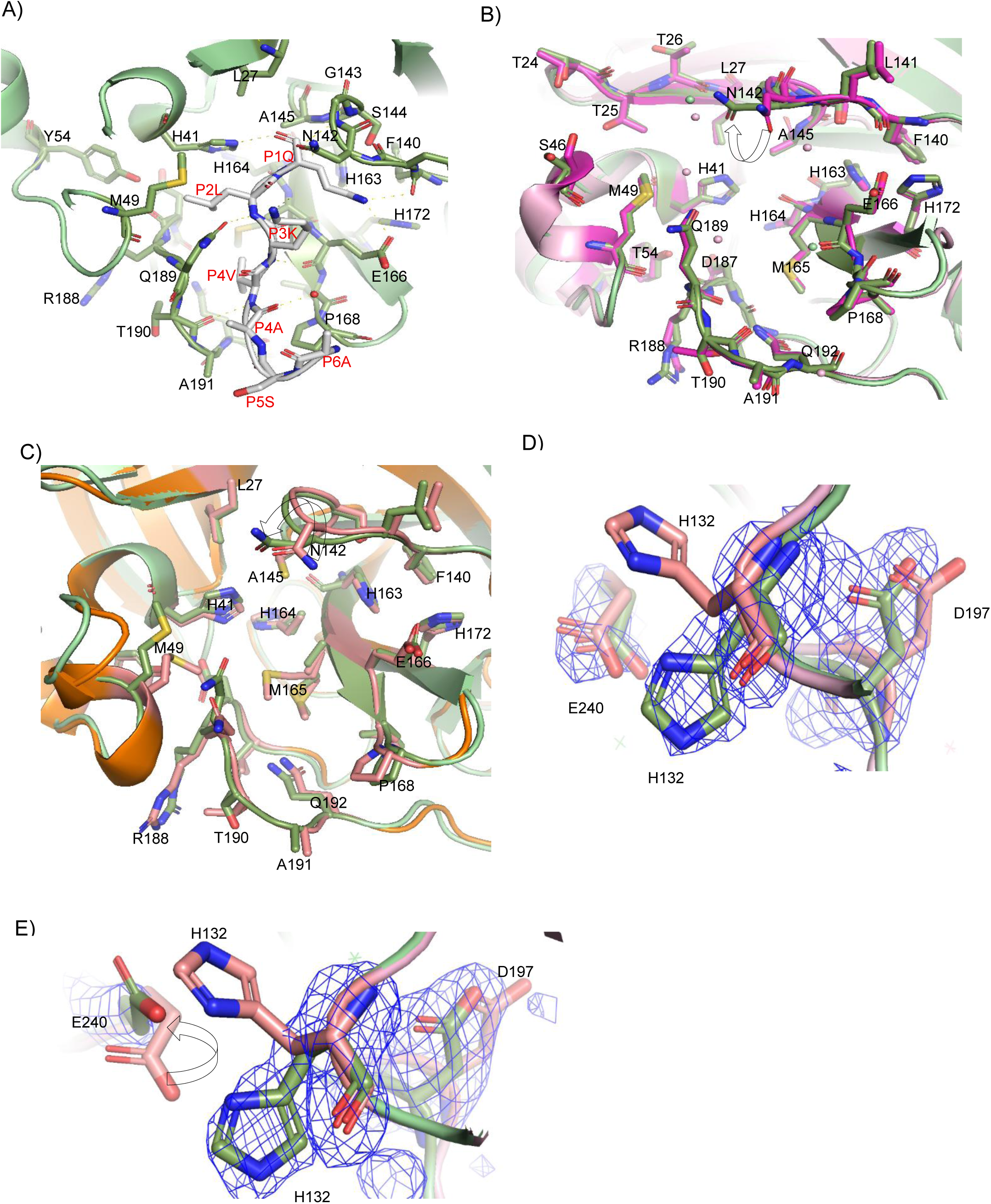
Recognition of Nsp8-Nsp9 substrate by Omicron 3CL-Mpro (11MJ). (A) substrate in the 3CL-Mpro Cys145Ala active site. Key Mpro residues mentioned in the text are labeled. Hydrogen bonds enabling 3CL-Mpro in recognition of substrate are shown as white dashed lines. (B) alignment of co-crystal structures of Omicron-3CL-Mpro and Nsp8-Nsp9 substrate (shown in pale green) with WT-3CL-Mpro and Nsp8-Nsp9 substrate (shown in light pink). (C) alignment of co-crystal structures of Omicron-3CL-Mpro and Nsp8-Nsp9 substrate (shown in pale green) with Apo Omicron-3CL-Mpro (shown in pale orange) (D) comparison of stacking orientation of His132 with Glu240 in presence (shown in pale green) and absence (shown in pale orange) of substrate. (E) comparison of edge to edge orientation of His132 with Glu240 in presence (shown in pale green) and absence (shown in pale orange) of substrate from another monomer. Electron density (Fo-Fc map) was carved at 3σ.

Nine hydrogen bonds were identified between 3CLpro (P132H) and the substrate. Seven backbone contacts were also observed. G143 and A145 main chain amides formed the oxyanion hole by donating hydrogen bonds to the P1 carbonyl oxygen, consistent with stabilization of the tetrahedral intermediate. H163, E166, and the main chain carbonyls of F140 and H164 formed hydrogen bonds with the P1Q side chain. Q189 formed a hydrogen bond with the P2L backbone. The backbone of P3K formed hydrogen bonds with the backbone groups of E166. The backbone nitrogen of P4V formed a hydrogen bond with the carbonyl group of T190. N142 interacted with P3K through an active-site water molecule.

The Nsp8-Nsp9 peptide bends away from the enzyme relative to the WT structure. Comparison with the WT co-crystal structure (7MGR) showed that the overall architecture of the active site remained conserved except for N142, which rotated approximately 90 degrees relative to WT-3CLpro and apo-Omicron 3CLpro (8R19) (Fig. 2B,C).

Comparison with apo-Omicron 3CLpro revealed changes near H132. In the apo structure, H132 engages in a stacking interaction with E240. In the substrate-bound structure, H132 rotates approximately 63 degrees in one monomer, forming an altered stacking interaction with E240. In the second monomer, both H132 and E240 rotate approximately 90 degrees to adopt an edge-to-edge configuration that supports a hydrogen bond. This rearrangement alters local interactions around residue 132. N197 also shifts by 1.4 Å toward H132 to form a hydrogen bond with the backbone nitrogen. An active-site water molecule is positioned to hydrogen bond with both the side chain and backbone of H132 (Fig. 2D, E; Sup Fig1). Whether the difference between protomers reflects a functional asymmetry or the distinct crystal-packing environment of each monomer remains to be established.

Together, these structural data show that the localized rearrangements remain confined to the vicinity of residue 132 and do not propagate to the substrate-binding pocket or catalytic residues, thereby preserving the structural integrity of the active site and providing a molecular basis for the WT-like catalytic efficiency observed in kinetic analyses.

### pH-Dependent Activity and Thermal Stability of WT and Variant 3CLpro Enzymes

To determine whether variant-associated substitutions alter physicochemical properties of 3CLpro beyond steady-state turnover, we next examined the pH dependence of catalytic activity and the thermal stability of WT and variant enzymes. These analyses were performed to assess whether distal mutations influence catalytic ionization behavior or overall structural stability despite the preserved active-site kinetics observed under standard assay conditions.

We determined the pH-activity profiles of the 3CLpro variants between pH 3.0 and pH 10.0. All enzymes showed maximal activity at approximately pH 7.5, consistent with the established ionization behavior of the C145 and H41 catalytic dyad (Fig. 4)^23,24^ The activity profiles were similar across WT and variant enzymes. The activity curves revealed two apparent pKa values for all enzymes (pKa1 ≈ 5.8 and pKa2 ≈ 8.2), indicating two ionization transitions affecting catalytic activity. No substantial differences in pH dependence were observed among the tested variants.

**Figure 3:**
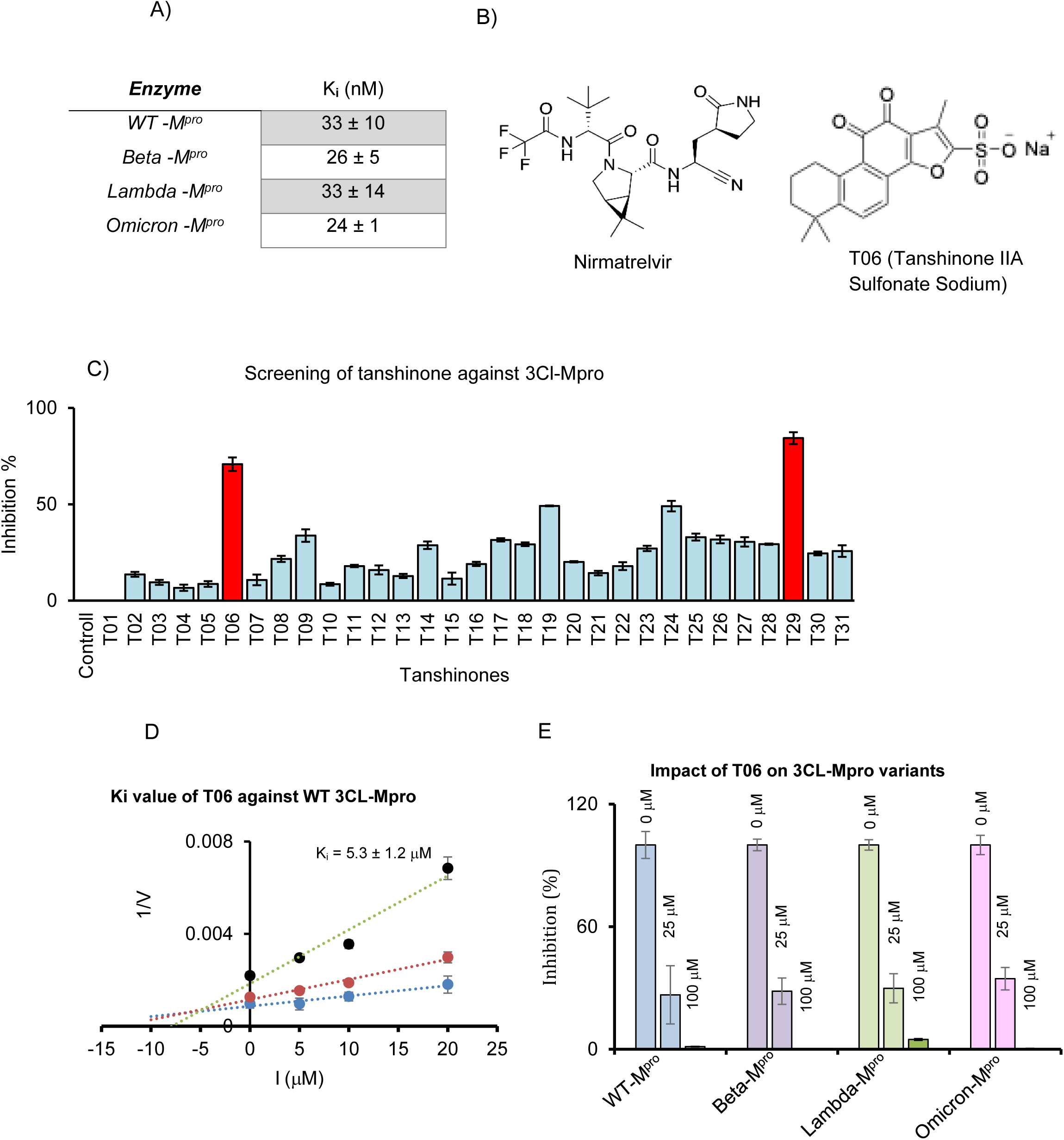
In vitro potency of Nirmatrelvir and Tanshinones for inhibiting the SARS-CoV-2 mutant Mpro activity in a plate-based FRET assay. A) Table of inhibition constants **(K_i_)** for Nirmatrelvir targeting 3CL-Mpro. 3CL-Mpro activity is monitored in the presence of increasing concentrations of Nirmatrelvir. B) Chemical structure of the inhibitor Nirmatrelvir. (C) screening of 31 tanshinones against WT-3CL-Mpro. (D) Determination of Inhibition constants (K_i_) for T06 targeting 3CL-Mpro. 3CL-Mpro activity is monitored in the presence of increasing concentrations of T06. (E) screening of T06 against mutant 3CL-Mpros

**Figure 4:**
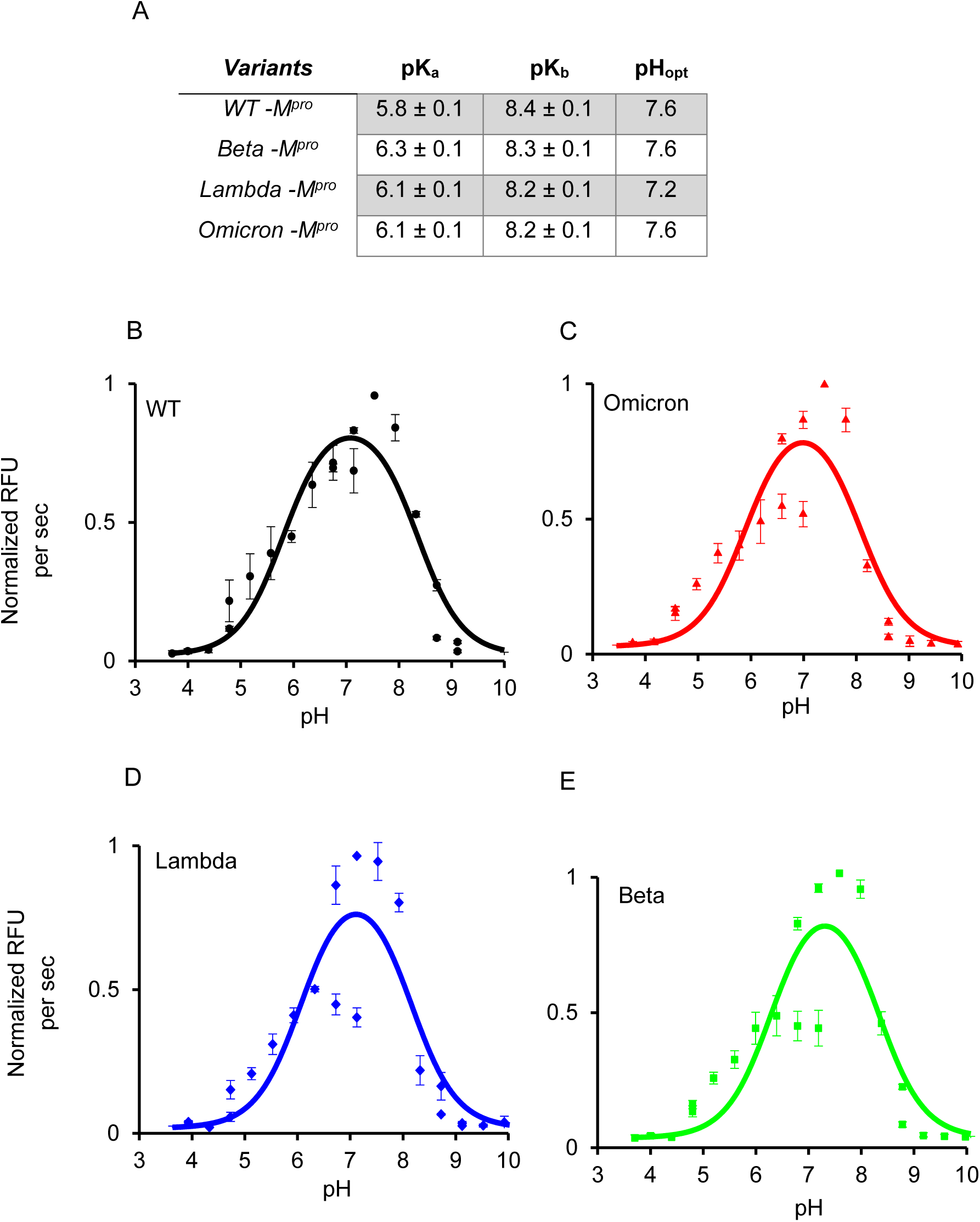
The effect of pH on catalytic activity of 3CL-Mpro from various variants of SARS-CoV-2. A) Table summarizing the ionization properties (pK_a_, pK_b_) and optimal catalytic pH (pH_opt_) of 3CL-Mpro from various variants of SARS-CoV-2. B-E) pH-dependent activity profile of WT, Omicron, Lambda, Beta 3CL-Mpro showing a bell-shaped curve. Initial rates were measured at a pH range of 3.5–10.

To assess whether mutations influence enzyme stability at elevated temperature, we examined the thermal resilience of WT and variant 3CLpro proteins. Enzymes were incubated at three temperatures, and residual activity was measured over time. At 25°C and 37°C, all variants-maintained activity for 96 hours without substantial loss (Fig. 5A,B). At 50°C, reduced stability of the Omicron variant was observed. After 1 hour at 50°C, the Omicron variant retained only 30 percent of its initial activity, indicating reduced thermal stability (Fig. 5C). In contrast, WT, Beta, and Lambda enzymes retained most of their initial activity under the same conditions. Reduced thermal stability of the P132H variant indicates that distal mutations can alter structural stability without impairing catalytic activity under physiological conditions^25,26^

**Figure 5:**
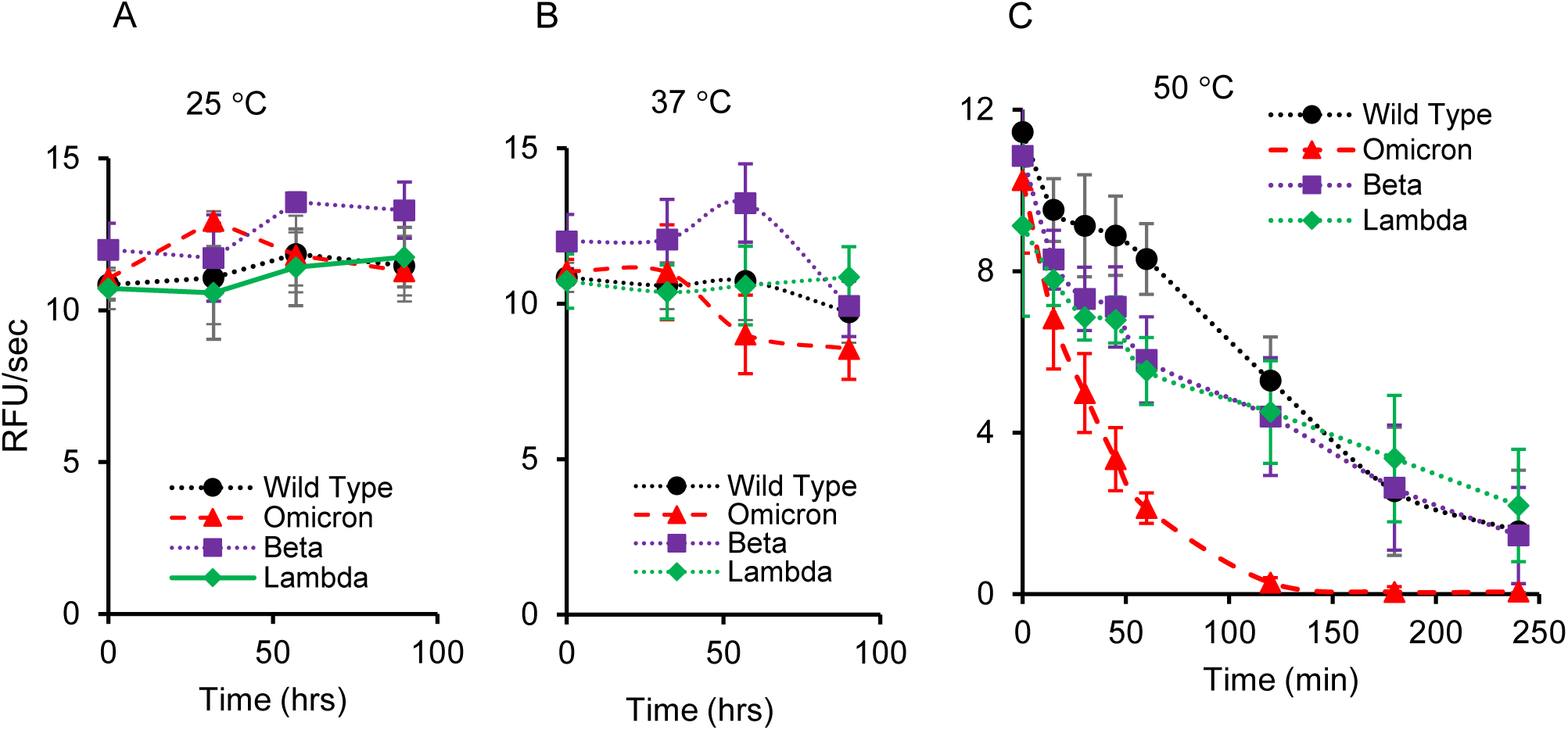
Temperature profiles of 3CL-Mpro from various variants of SARS-CoV-2. The temperature-activity profile was measured at three temperature - 25°C (A), 37°C (B), 50°C (C) in 50 mM Tris–HCl buffer (pH 7.5).

Together, these findings indicate that the P132H substitution selectively reduces the thermal stability of 3CLpro under heat stress while preserving its catalytic ionization profile and enzymatic activity under physiological conditions, showing that distal substitution can therefore weaken the fold while leaving catalysis intact.

### Interaction of Nirmatrelvir with 3CLpro variants

To determine whether variant-associated substitutions alter the susceptibility of 3CL-Mpro to clinically relevant protease inhibition despite preserved catalytic activity, we next assessed the inhibitory potency of Nirmatrelvir (PF-07321332) against the Omicron, Beta, and Lambda variants using a fluorescence-based inhibition assay. The inhibition constant (K_i_) for WT 3CLpro was 33 ± 10 nM, which was similar to the values obtained for the variants (Omicron K_i_ = 24 ± 1 nM; Beta K_i_ = 26 ± 5 nM; Lambda K_i_ = 33 ± 14 nM) (Fig. 3A). No substantial differences in Ki values were observed among WT and variant enzymes^10,11,12^.

Together, these data indicate that the examined distal substitutions do not measurably alter Nirmatrelvir binding, supporting the retained inhibitor susceptibility of major SARS-CoV-2 variants and the durability of 3CLpro-targeted antiviral therapy.

### Screening of Tanshinones against 3CLpro

Given the preserved susceptibility of all tested variants to Nirmatrelvir, we next asked whether 3CLpro could also be inhibited by alternative small-molecule scaffolds. Based on prior reports implicating tanshinone derivatives as coronavirus protease inhibitors^27^ and human cystein cathepsins protease inhibitors^20,21^, we screened a panel of 31 tanshinones using a plate-based fluorescence assay with a peptide substrate corresponding to the Nsp4-Nsp5 cleavage site. Several compounds inhibited WT 3CLpro, with the most active candidate, T06 (Tanshinone IIA Sulfonate Sodium), reducing enzymatic activity by approximately 70% at 50 µM (Fig. 3.C). Kinetic analysis yielded an inhibition constant (Ki) of 5.3 ± 1 µM for WT 3CLpro (Fig. 3.D). T06 was subsequently tested against variant enzymes, including Omicron, Beta, and Lambda 3CLpro, and inhibited all tested variants under the same assay conditions. These results indicate that T06 inhibits both WT and variant 3CLpro with comparable potency under the conditions tested.

## Discussion

Our biochemical and structural analyses show that mutations distal to the active site do not disrupt the core catalytic machinery of SARS-CoV-2 3CLpro. Consistent with their location outside the catalytic dyad, the K90R, G15S, and P132H substitutions preserved catalytic efficiency, pH dependence, and susceptibility to Nirmatrelvir^10,11,23^. This conservation indicates that the fundamental chemical steps governed by Cys145 and His41 remain intact across these variants, consistent with the established catalytic mechanism of 3CLpro^28^. The persistence of a bell-shaped pH–activity profile further supports the conclusion that the acid–base equilibria underlying the catalytic cycle are unaffected by these substitutions^23^. This pattern contrasts with the principal routes to nirmatrelvir resistance, which involve substitutions at or adjacent to the active site, such as E166V, L50F, and S144A^29^; the distal substitutions examined here leave inhibitor binding intact, supporting a strategy that targets the conserved catalytic core.

The thermal stability differences observed for the Omicron 3CLpro suggest that this variant may possess distinct biophysical characteristics relative to the WT and other circulating variants. Although all proteases retained more than 95% activity over 100 hours at physiological temperatures, the Omicron enzyme showed a marked reduction in stability under elevated-temperature stress. This enhanced thermal lability may indicate that the Omicron protease occupies a narrower stability margin, potentially rendering its fold more sensitive to perturbation. One plausible contributor is the P132H substitution, which is absent from the other variants evaluated and may introduce subtle destabilizing effects, either by perturbing the hydrophobic core or modifying domain-domain packing. Because all variants remained stable at physiological temperature, this reduced thermal resilience is unlikely to affect 3CLpro function during infection and instead provides a measurable biophysical signature of the P132H fold.^25,26^.

The central mechanistic question raised by Omicron P132H is how efficiently this enzyme can process cleavage sites beyond Nsp4-Nsp5, compared to WT-3CLpro. Both WT and P132H cleave this site with markedly lower efficiency than the canonical Nsp4-Nsp5 junction.^13^ Our kinetic measurements confirm that the reduced turnover of the Nsp8-Nsp9 junction is not a mutation-specific defect but an intrinsic property of the substrate. The structural preservation of the D289-R131 salt bridge and the overall domain arrangement in the Omicron enzyme further supports the conclusion that the mutation does not substantially alter substrate alignment or binding. Our crystallographic analysis indicates that in the substrate-bound state, H132 adopts an alternative conformation relative to the apo Omicron 3CLpro structure. This repositioning enables the formation of new interactions with E240 and adjacent residues that effectively rebuild the local hydrogen-bonding network. These observations indicate substrate-induced reorganization of the surrounding environment of H132. Because the wild-type and P132H enzymes process these substrates with comparable efficiency, the mutation is unlikely to alter viral polyprotein processing or pathogenicity.

The initial screening of 31 tanshinones revealed T06 as the lead candidate against the WT-3CLpro. Tanshinones, natural diterpenoids derived from *Salvia miltiorrhiza*, are known for their polycyclic aromatic structures. T06 inhibited WT 3CLpro and retained inhibitory activity against the tested variant enzymes. T06 therefore provides a preliminary lead for further study, although its mechanism of inhibition and selectivity remain to be established before its potential as a pan-coronaviral scaffold can be assessed.

A broader implication of these findings is that 3CLpro possesses substantial structural plasticity^30^. Mutations at positions that may contribute to scaffold instability, can be buffered by substrate-induced loop stabilization, allowing the enzyme to reorganize local interactions in ways that maintain the integrity of the catalytic environment^31^. These properties can be an explanation how circulating variants were able to accumulate substitution while retaining full enzyme competence. It also points to a challenge for antiviral design. Inhibitors that rely on rigid interactions with regions capable of adopting alternative conformations may be less resilient to future sequence variation, whereas molecules that engage conserved elements of the catalytic machinery are more likely to retain potency. Several limitations temper these conclusions. We examined two of the eleven polyprotein junctions and threesingle-substitution variants; effects at the remaining cleavage sites, in variants carrying multiple co-occurring substitutions, and during cellular infection remain to be tested.

Our study indicates a structural mechanism through which 3CLpro may be able to preserves catalytic function despite having mutations that alter local stability^25,26^. The ability of the substrate to stabilize alternative H132 conformations provide a hypothesis for the resilience of the protease during viral evolution. This compensatory mode of mutation tolerance must be considered in future efforts to develop inhibitors capable of withstanding ongoing diversification of SARS-CoV-2.

### Conclusion

SARS-CoV-2 3CLpro tolerates the distal substitutions carried by the Beta (K90R), Lambda (G15S), and Omicron (P132H) variants. All four enzymes processed both the Nsp4-Nsp5 and Nsp8-Nsp9 substrates with comparable catalytic efficiency and displayed near-identical pH-rate profiles, indicating conserved substrate recognition and catalytic mechanism. The crystal structure of Omicron 3CLpro bound to an Nsp8-Nsp9 peptide explains this resilience: the fold and active-site geometry are unperturbed, with the P132H side chain locally rebuilding its contacts in a substrate-dependent manner. The main consequence of P132H is thermal rather than catalytic. These results support 3CLpro as a durable antiviral target, and our tanshinone screen extends the search for drugs complementing covalent inhibitors such as Nirmatrelvir.

### Materials

Site-directed mutagenesis Q5 kit from NEB (E0554S), (MCA)`AVLQSGFR`[`K`(Dnp)]`RR` was synthesized (>95% purity) by GenScript, Piscataway, USA, FRET-based peptides, (DABCYL)KRVVKLQNNEIME(EDANS)^13^ and LRANSAVKLQNNELSPVALR was custom synthesized (>98% purity) by Biomatik Co., Ontario, Canada, Amicon^®^ Ultra - 15 Centrifugal Filters were from Merck Millipore Ltd, Ireland), 3CLpro inhibitor Nirmatrelvir from A2B Chem, LLC (San Diego, CA). Sodium tanshinone IIA sulfonate (Carbosynth Ltd. UK).

## Methods

### Cloning and Protein Expression

The gene encoding full-length WT-SARS-CoV-2 3CLpro (NCBI Reference Sequence: NC_045512.2), containing an N-terminal methionine removed co-translationally by endogenous methionine aminopeptidase and C-terminal FLAG, Myc, and His tags, was provided by Dr. Christopher Overall (University of British Columbia). A PreScission protease cleavage site was introduced using site-directed mutagenesis, and the resulting construct was used as the template for generating all 3CLpro variants. All plasmids were sequence-verified by Sanger sequencing. His₆-tagged WT 3CLpro was expressed in *Escherichia coli* BL21(DE3), and all 3CLpro mutants, including the catalytically inactive C145A variant, as well as PreScission protease, were expressed in *E. coli* BL21 Star(DE3). Cultures (2-6 L) were grown in Lysogeny Broth (LB) at 37°C with 100 µg/mL ampicillin to an OD₆₀₀ of 0.6, cooled to 16°C, and induced with 0.5 mM IPTG overnight. Cells were harvested by centrifugation at 5,000 × g for 10 min at 15°C (Avanti J-E centrifuge, Beckman Coulter) and resuspended in lysis buffer (50 mM Tris, pH 7.5, 300 mM NaCl).

### Protein Purification

Cells were lysed by sonication on ice and clarified by centrifugation at 20,000 × g for 30 min at 4°C. The supernatant was applied to Ni-NTA resin (QIAGEN, USA) equilibrated in lysis buffer. Bound proteins were eluted with 50 mM Tris (pH 7.5), 300 mM NaCl, and 300 mM imidazole. Eluted 3CLpro proteins were buffer-exchanged overnight in the presence of PreScission protease to remove imidazole and cleave the His tag. The sample was passed again over Ni-NTA to remove non-cleaved protein and protease. The flow-through was loaded onto a Superdex 200 size-exclusion column (Cytiva Life Sciences, Uppsala/Sweden) equilibrated with 50 mM Tris (pH 7.5), 150 mM NaCl using an ÄKTA Purifier (Cytiva Life Sciences, Uppsala/Sweden). Purified protein was concentrated and stored at −80°C. PreScission protease was purified identically and stored at −80°C in buffer containing 5 mM β-mercaptoethanol.

### Enzymatic Activity Assays

Protease activity was measured at 23°C using the fluorogenic substrates (MCA)`AVLQ`↓`SGFR`[`K`(Dnp)]`RR` corresponding to the Nsp4-Nsp5 cleavage site, and (DABCYL)`KRVVKLQ`↓`NNEIME`(EDANS)^32^ corresponding to the Nsp8-Nsp9 site. Assay with `LQ`↓`SG` was performed in 50 mM Tris (pH 7.5) containing 10 mM DTT. Fluorescence was monitored on a BioTek Synergy H4 microplate reader (Santa Clara, USA; λ_ex_ = 320 nm, λ_em_ = 405 nm). For steady-state kinetics, 3CLpro was used at 100 nM, and substrate concentrations ranged from 0 to 60 µM. Initial velocities (first 2 min) were converted to product concentration using an MCA calibration curve. Assay with `LQ`↓`NN` was also performed in 50 mM Tris (pH 7.5) containing 10 mM DTT. Fluorescence was monitored on a BioTek Synergy H4 microplate reader (λ_ex_ = 336 nm, λ_em_ = 490 nm respectively). For steady-state kinetics, 3CLpro was used at 500 nM, and substrate concentrations ranged from 0 to 80 µM. Initial velocities (first 2 min) were converted to product concentration using an EDANS calibration curve. Kinetic parameters (K_M_ and k_cat_) were obtained by nonlinear regression using the Michaelis-Menten equation in using GraphPad Prism (Version 5.0, GraphPad Software, La Jolla, California, USA). All assays were performed in two independent experiments, each in triplicate (2 biological replicates x 3 technical).

### Inhibition Assays

Inhibition of 3CLpro was assessed using substrate concentrations ranging from 4 to 16 µM and inhibitor concentrations ranging from 0 to 100 nM. Reactions were carried out at 23°C in 50 mM Tris (pH 7.5) containing 10 mM DTT. Fluorescence was monitored for 2 min at λ_ex_ = 320 nm and λ_em_ = 405 nm using a BioTek Synergy H4 microplate reader.

Initial velocities were calculated from the linear portion of each fluorescence trace. Activity in the absence of inhibitor was defined as 100 percent, and relative activity at each inhibitor and substrate concentration was plotted as initial velocity (v) versus inhibitor concentration [I]. Dixon plots were generated by plotting 1/v against [I] at multiple substrate concentrations. Linear regression was performed for each line, and the intersection point of the regression lines provided the apparent inhibition constant (K_i_). All regression analyses and Dixon plot fitting were performed using GraphPad Prism 5 (Version 5.0, GraphPad Software, La Jolla, California, USA). All assays were performed in two independent experiments, each in triplicate.

### pH and Temperature Profiling

pH dependence of WT and mutant 3CLpro was measured at 100 nM enzyme and 10 µM substrate in buffers spanning pH 3-10 (acetate, phosphate, Tris, borate, CHES), all containing 10 mM DTT. pK_a_ values were calculated using SigmaPlot (Grafiti LLC, USA). Temperature dependence was assessed at 25°C, 37°C, and 50°C with 100 nM enzyme and 10 µM substrate.

### Co-Crystallization of Peptide-Bound Omicron 3CLpro

Peptide (Nsp8-Nsp9 sequence; LRANSAVKLQNNELSPVALR) was dissolved at ∼1 mg/mL in 0.1 M HEPES (pH 8.0) containing 4% DMSO and mixed with Omicron 3CLpro (C145A; 6 mg/mL in 50 mM Tris-HCl, 150 mM NaCl) to a final peptide concentration of 2 mM. Crystals were obtained by sitting-drop vapor diffusion at 22°C by mixing 0.7 µL of protein– peptide solution with 0.7 µL reservoir solution (25% PEG 3350, 0.1 M BT (pH 5.5), 0.2 M Ammonium sulfate). Crystals formed within 3 weeks and were transferred into reservoir solution supplemented with cryoprotectant (25% ethylene glycol) and 1 mM peptide before flash-freezing in liquid nitrogen. Diffraction data were collected at 100 K using a DECTRIS EIGER2 XE 16M detector at beamline BL12-2, Stanford Synchrotron Radiation Lightsource (SSRL), with λ = 0.98 Å. Data were processed with iMOSFLM v7.2.1 and scaled with SCALA^33,34^. Molecular replacement was performed using WT 3CLpro (PDB 7JP1)^15^. Models were refined in PHENIX and manually adjusted in COOT^35,36^. Model quality was verified using MolProbity ^37^. Structural figures were prepared with PyMOL (Schrödinger LLC). PDB id of the structure is 11MJ

### Inhibition Assays with Tanshinones

For fluorogenic peptide substrate hydrolysis, 100 nM WT-3CLpro was added with or without 50 μM tanshinones in 100 μL, 50 mM Tris buffer, pH 7.5, containing 10 mM DTT, and 10 μM fluorogenic peptide substrate based on native Nsp4-Nsp5 cleavage site. Then best candidate was screened against other 3CLpro variants of SARS-CoV2. For K_i_ value determination, substrate concentrations used from 8 to 25 µM and inhibitor concentrations used from 0 to 50 μM. Reactions were carried out at 23°C in 50 mM Tris (pH 7.5) containing 10 mM DTT. Fluorescence was monitored for 2 min at λ_ex_ = 320 nm and λ_em_ = 405 nm using a BioTek Synergy H4 microplate reader.

Initial velocities were calculated from the linear portion of each fluorescence trace. Activity in the absence of inhibitor was defined as 100 percent, and relative activity at each inhibitor and substrate concentration was plotted as initial velocity (v) versus inhibitor concentration [I]. Dixon plots were generated by plotting 1/v against [I] at multiple substrate concentrations. Linear regression was performed for each line, and the intersection point of the regression lines provided the apparent inhibition constant (K_i_).

## Author contributions

DS, SYTD, and DB designed the study, validated results, and were responsible for the administration of project. DS, SYTD, EM performed substrate kinetics, Ki determination. DS was responsible together with FvP and YSC for the crystallization and structural analysis of the peptide-Omicron3CL-Mpro complex. DS and SYTD produced and purified SARS-CoV-2 3CL-Mpro along with C145A mutant. DS, SYTD, and DB wrote the original draft of the manuscript. All authors reviewed and revised the final manuscript. DB acquired funding for the study.

## Acknowledgements

This work was supported by Canadian Institutes of Health Research grants (PJT-155979, CPG-158275), Collaborative Health Research Project (NSERC CHRP 523434-18) and the NSERC discovery grant (LJGP GR003266). DB was supported by Canada Research Chair award funding.

## Notes

### Competing Interest Statement

The authors have declared no competing interest.

### Summary of Updates

the manuscript has been revised to update the claims and removed strong words as well. electron density map has beed added.

